# The eukaryotic ribosomal protein S15/uS19 is involved in fungal development and its C-terminal tail contributes to stop codon recognition

**DOI:** 10.1101/2020.02.09.940346

**Authors:** Tan-Trung Nguyen, Guillaume Stahl, Michelle Déquard-Chablat, Véronique Contamine, Sylvie Hermann-Le Denmat

## Abstract

S15/uS19 is one of the fifteen universally conserved ribosomal proteins of the small ribosomal subunit. While prokaryotic uS19 is located away from the mRNA decoding site, cross-linking studies identified eukaryotic uS19 C-terminal tail as contacting the A site on the 80S ribosome. Here, we study the effects of uS19 mutations isolated as translation accuracy mutations in the filamentous fungus *Podospora anserina*. All mutations alter residues of uS19 C-terminal tail, and cluster to the eukaryote-specific decapeptide 138-PGIGATHSSR-147. All mutations modify fungal development and cytosolic translation, albeit differently. Two mutations (P138S and S145F) increase fungus longevity and display mild effects on translation, while others (G139D and G139C) decrease longevity, have stronger effects on translation and confer hypersensitivity to paromomycin. By mimicking *P. anserina* mutations in the yeast *Saccharomyces cerevisiae RPS15* gene, we further show that P138S and S145F induce hyperaccurate recognition of the three stop codons, whereas G139D and G139C impair UAG and UAA codon recognition. Noteworthy, in *P. anserina*, uS19 genetically interacts with the eRF1 and eRF3 release factors. All together, our data indicate that uS19 C-terminal tail contributes *in vivo* to eukaryotic translation termination, and identify key amino acids of uS19 that potentially modulate eRF1-eRF3 interaction in the pre-termination complex.

**Graphical abstract:** 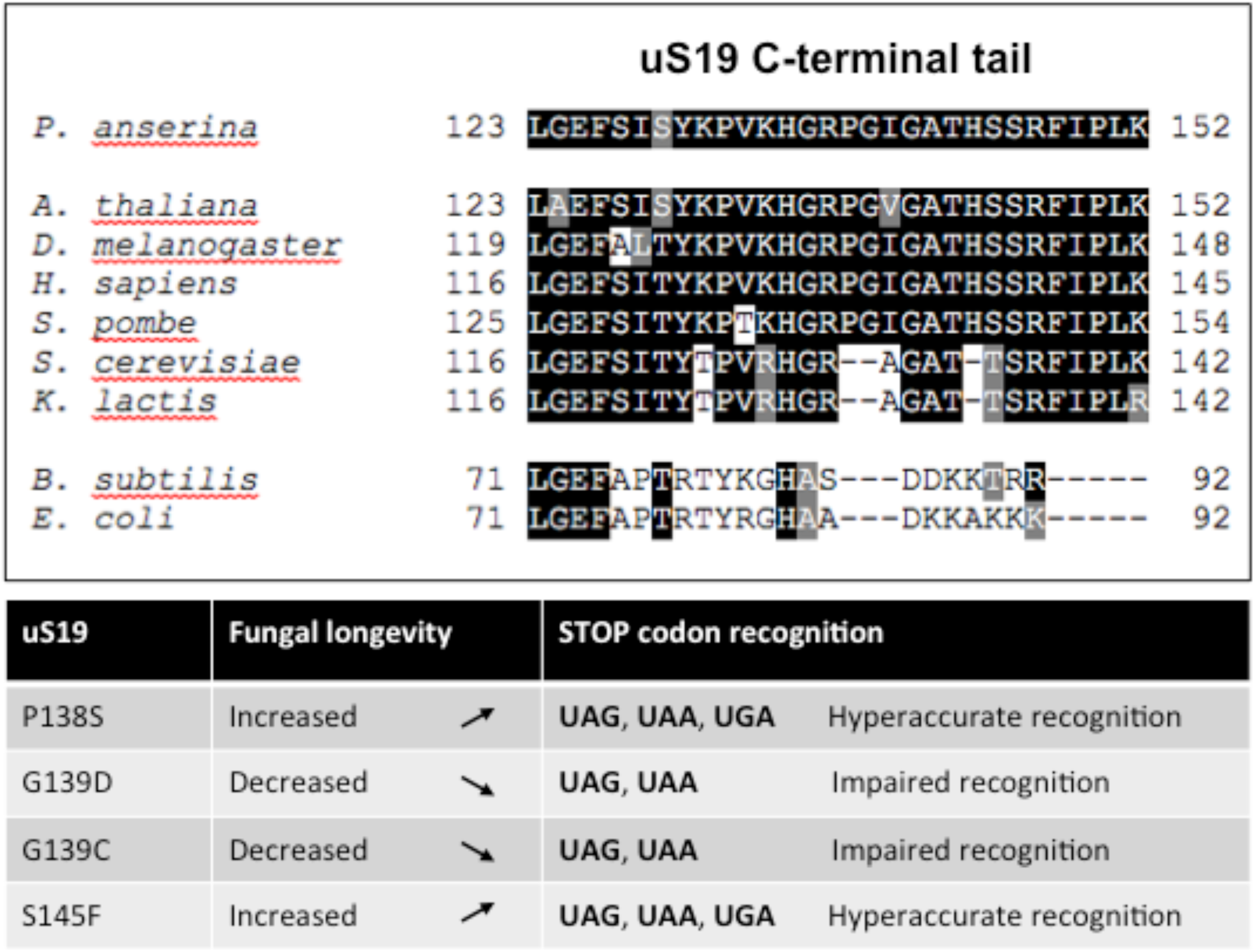

**Abbreviated Summary:** S15/uS19 is a conserved small ribosomal protein that in eukaryotes harbors a flexible C-terminal extension proposed to interact with the A site mRNA codon during translation. Here, we describe how C-terminal variants variously affect *Podospora anserina* development and longevity and impact fungal ribosome and polysome formation. We reveal that stop codon recognition is significantly altered by the presence of those C-terminal variants, which either expand or on the contrary restrict termination ambiguity.

## Introduction

In all organisms, ribosomes are responsible for protein synthesis. The small subunit is engaged in decoding the information encoded in mRNA, while peptide bond synthesis occurs at the large ribosomal subunit. In eukaryotes, the cytoplasmic large 60S and small 40S subunits are together composed of 80 distinct proteins and four ribosomal RNA species. The 60S subunit is assembled from 25S-28S rRNA, 5.8S rRNA, 5S rRNA and 47 proteins, of which seven are specific to eukaryotes, 20 to eukaryotes and archea, and 20 are found in all kingdoms (Lecompte *et al*., 2002). Similarly, the 40S subunit consists of 18S rRNA and 33 proteins, of which five are specific to eukaryotes, 13 to eukaryotes and archea, and 15 are found in all kingdoms (Lecompte *et al*., 2002; Rabl *et al*., 2011; Ban *et al*., 2014).

Ribosomal proteins (r-proteins), in addition to being constituents of ribosomes, play major roles in rRNA processing, ribosome assembly, nucleo-cytoplasmic transport of ribosomal subunits, and in the functioning of the translational machinery itself (Brodersen and Nissen, 2005; Ferreira-Cerca *et al*., 2005; Wilson and Nierhaus, 2005; Pöll *et al*., 2009; Woolford and Baserga, 2013; Graifer and Karpova, 2015). The assignment of exact ribosomal functions to individual r-proteins is not easy to achieve considering the highly cooperative nature of protein-protein and protein-RNA interactions in the ribosome. Whereas the roles of a number of bacterial r-proteins in translation have been understood (Brodersen and Nissen, 2005; Wilson and Nierhaus, 2005; for reviews), the contributions eukaryotic r-proteins make to translation have been less addressed, and the simple idea that eukaryotic r-proteins share the same functional properties in ribosomes as their bacterial counterparts can be erroneous and is a subject of debate (see Graifer and Karpova, 2015 and references therein).

What adds to the difficulty is that r-proteins have N and C-terminal extensions that are specific to eukaryotes (Rabl *et al*., 2011), and these could participate in eukaryote-specific functions of r-proteins. One potential example of this is uS19 (also known as S15, the ortholog of bacterial S19p; (Ban *et al*., 2014)). In yeast, this essential r-protein was shown to be involved in the nuclear export of 40S subunit precursors (Léger-Silvestre *et al*., 2004), a role conserved in mammalian cells (Rouquette *et al*., 2005), and it was suggested that the N-terminal extension of uS19 (see Fig. 1A) may be involved (Léger-Silvestre *et al*., 2004).

**Fig. 1.**
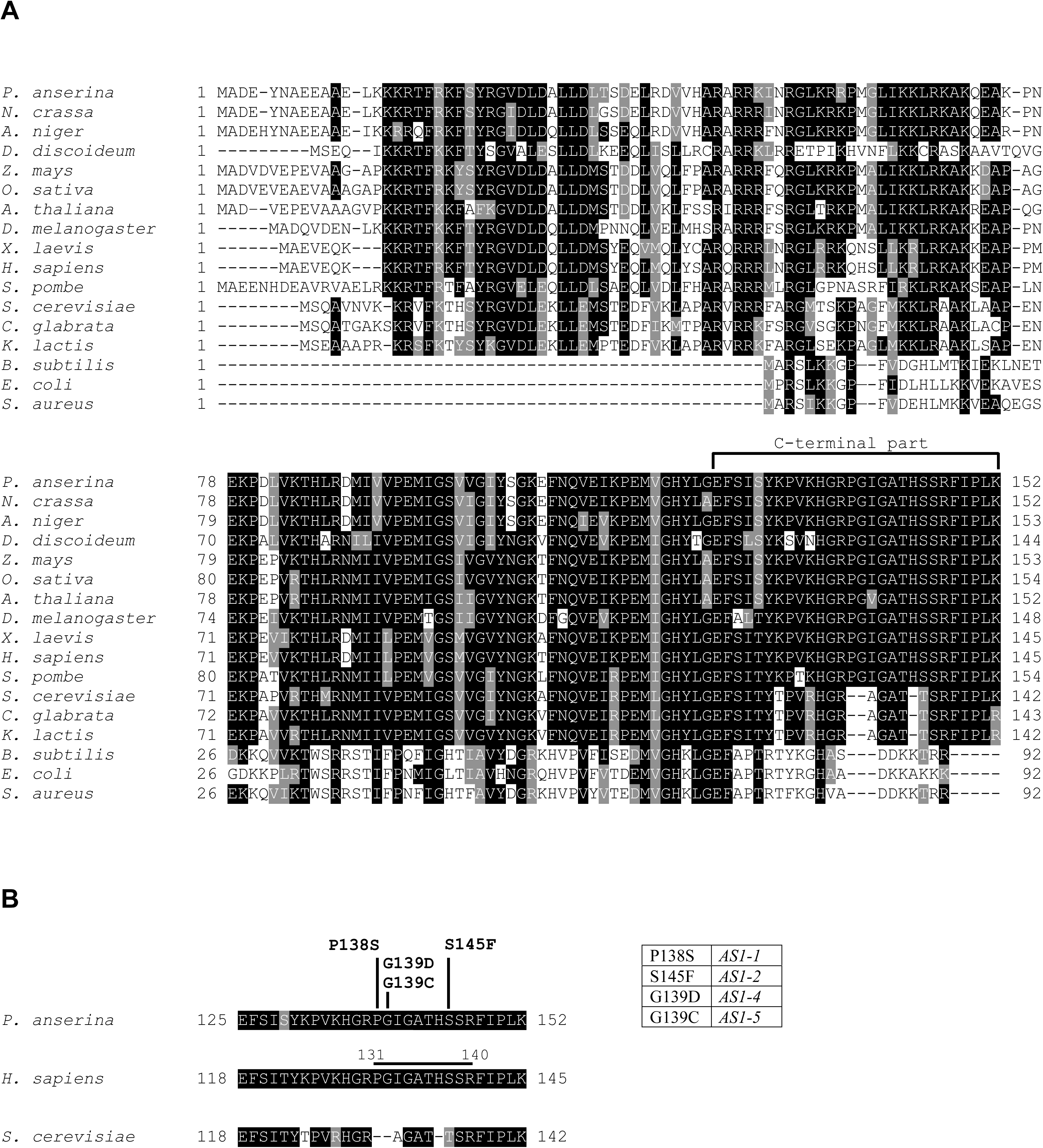
*P. anserina AS1* mutations affect nearby residues of the highly conserved C-terminal part of the eukaryotic r-protein uS19. A. Multiple sequence alignment of full-length uS19 protein from fourteen eukaryotic and three prokaryotic organisms. Alignment was carried out using ClustalW2 program (Larkin *et al*., 2007). Conserved amino acids (aas) are boxed in black (identical) and gray (similar). Numbering refers to full-length protein sequence and amino acid sequences were retrieved from the NCBI database (http://www.ncbi.nlm.nih.gov). NCBI references are (order is as in the alignment): *Podospora anserina* XP_001907110, *Neurospora crassa* XP_965164, *Aspergillus niger* XP_001397802, *Schizosaccharomyces pombe* NP_594357, *Dictyostelium discoideum* XP_638126, *Zea mays* NP_001147395, *Oryza sativa* NP_001051572, *Arabidopsis thaliana* (isoform 1, which one shares the highest sequence identity with *P. anserina* uS19 polypeptide), NP_171923, *Drosophila melanogaster* NP_611136, *Xenopus laevis* NP_001089043, *Homo sapiens* NP_001009, *Saccharomyces cerevisiae* NP_014602, *Candida glabrata* XP_446019.1, *Kluyveromyces lactis* XP_455435, *Bacillus subtilis* NP_388001, *Escherichia coli* NP_289877 and *Staphylococcus aureus* YP_005747034. B. C-terminal part of *P. anserina*, *H. sapiens* and *S. cerevisiae* uS19 r-proteins as indicated in A. Amino acid changes induced by *P. anserina AS1* mutations are indicated. Match between the *AS1* alleles and residue substitution is shown in the inserted table. In yeast mimetic mutants (Fig. 4 and 5), the 25 aas C-terminal end of *S. cerevisiae* uS19 has been exchanged for the 28 aas C-terminal end of *P. anserina* r-protein. The decapeptide 131-PGIGATHSSR-140 (*H. sapiens* numbering) mapped to neighbor the A site mRNA codon in human translating ribosome (Khairulina *et al*., 2010) is pointed out by a line.

Eukaryotic uS19 localizes to the head domain of the small ribosomal subunit, a location conserved with bacteria (Ferreira-Cerca *et al*., 2007; Rabl *et al*., 2011). Interestingly, while the bacterial polypeptide is positioned away from the decoding site (Brodersen and Nissen, 2005) eukaryotic uS19 was found to neighbor it and to contact the mRNA during the translation process (Bulygin *et al*., 2002). Furthermore, site-directed cross-linking studies performed with diverse mammalian reconstituted ribosomal complexes showed that uS19 is one of the only, if not the only, eukaryotic r-proteins that interacts with the A site mRNA codon during the initiation, elongation and termination steps of translation (Bulygin *et al*., 2005; Pisarev *et al*., 2006; Pisarev *et al*., 2008). Cross-link was mapped in the C-terminal tail of human uS19 (Khaĭrulina *et al*., 2008), most probably in the eukaryote-specific peptide PGIGATHSSR (Khairulina *et al*., 2010) (see Fig. 1A and B). However, the lack of structural resolution of the C-terminal sequence of uS19 (Rabl *et al*., 2011; Lomakin and Steitz, 2013) prevents visualization of uS19 interaction with the mRNA at the decoding site that would confirm the documented *in vitro* cross-linking data. This, in turn, makes establishing the functional role of uS19 in eukaryotic translation difficult. Nevertheless, hypothetic roles of the PGIGATHSSR decapeptide have been proposed, including selection of the initiation codon during initiation (Pisarev *et al*., 2008), stabilization of the mRNA codon at the decoding site during elongation (Khaĭrulina *et al*., 2008; Khairulina *et al*., 2010), as well as interaction with the polypeptide chain release factor eRF1 during termination (Khairulina *et al*., 2010; Graifer and Karpova, 2012; Graifer and Karpova, 2015, for reviews).

To study the ribosomal control of translation fidelity in the filamentous fungus *Podospora anserina*, Marguerite Picard’s group has carried out in the 70’s very elegant two-step genetic screens that recovered, among other ribosomal mutations, mutations in the *AS1* gene that codes for uS19 (Picard, 1973; Picard-Bennoun, 1976; Picard-Bennoun, 1981; Dequard-Chablat and Sellem, 1994). We report here that the four *AS1* mutations identified in those studies all alter residues of the PGIGATHSSR decapeptide that is identical between the *P. anserina* and human r-proteins (Fig. 1B). Interestingly, these *AS1* mutations antagonize the effect of missense mutations in the genes encoding the release factors eRF1 and eRF3 (Picard, 1973; Picard-Bennoun, 1976; Gagny and Silar, 1998; This work). By dissecting the phenotypic traits associated with each *AS1* mutation in *P. anserina* and studying the translation defects associated with each *AS1* mutation in *P. anserina* and in yeast, we demonstrate the importance of uS19 to fungal development and the critical role uS19 C-terminal tail plays in stop codon recognition. Our *in vivo* findings confirm for the first time hypotheses formulated to explain *in vitro* cross-linking results that had identified contacts of the uS19 C-terminal tail with the A site mRNA codon (Graifer and Karpova, 2015).

## Results

### AS1 anti-suppressor mutations alter residues of uS19 C-terminal tail

Mutated alleles of the *P. anserina AS1* gene were obtained in the course of two-step genetic screens aimed to study, four decades ago, the ribosomal control of translational ambiguity in eukaryotes (Picard, 1973; Picard-Bennoun, 1976; Picard-Bennoun, 1981). First, many *su* mutations acting as informational <u>su</u>ppressors (*i.e.*, they increased translational error) have been obtained, and secondly, starting from some *su* mutations, <u>a</u>nti-<u>s</u>uppressor mutations (*AS*) antagonizing the effect of *su* mutations have been isolated (Picard-Bennoun, 1976; Coppin-Raynal, 1981; Coppin-Raynal *et al*., 1988, for reviews). One of these genetic screens was based on an ascospore defective color phenotype associated with the mutation *193* suspected at the time to be a nonsense mutation. Indeed, sequencing revealed mutation *193* as being an UGA stop codon in the open reading frame encoding a polyketide synthase involved in melanin formation (Coppin and Silar, 2007). Phenotypically, while the stop-codon mutation *193* affected ascospore pigmentation, in a *193 su* double mutant context, pigmentation was restored (partially or totally), and in a *193 su AS* triple mutant context, ascospore pigmentation was anew defective (see Table 1). Some of the *su* and *AS* genes have been cloned (Debuchy and Brygoo, 1985; Silar and Picard, 1994; Silar *et al*., 1997; Gagny and Silar, 1998; Dequard-Chablat and Silar, 2006), including the *AS1* gene that codes for the cytosolic r-protein uS19 (Dequard-Chablat and Sellem, 1994; therein uS19 was named S12 according to electrophoretic nomenclature). In all cases, only actors of the translation machinery were identified as initially predicted by the elegant pioneer genetic approaches carried out by Marguerite Picard (Picard, 1973; Picard-Bennoun, 1976; Picard-Bennoun, 1981).

**Table 1.** Selection mode of the *AS1* mutant alleles.

| *AS1* allele | Genetic context of the selection<sup>a</sup> | Criteria of selection | Missense mutation in *AS1*<sup>b</sup> | Codon substitution | Reference |
| --- | --- | --- | --- | --- | --- |
| *AS1-1* | *193 su1-42* | Ascospore color | **CCA**->**TCA** | P138S | Picard-Bennoun (1976); Belcour *et al.* (1991) |
| *AS1-2/AS1-3<sup>c</sup>* | *193 su1-1* | Ascospore color | **TCT**->**TTT** | S145F | Picard-Bennoun (1976) |
| *AS1-4* | *193 su1-1* | Ascospore color | **GGT**->**GAT** | G139D | Belcour *et al.* (1991); Déquard-Chablat and Sellem (1994) |
| *AS1-5* | *su2-5* | Improved growth | **GGT**->**TGT** | G139C | Déquard-Chablat and Sellem, (1994); M. Picard, unpubl. work |

**a.** The *193* mutation is a nonsense mutation in the polyketide synthase encoding gene (Coppin and Silar, 2007); *su1-1* and *su1-42* are mutant alleles of the *su1* gene encoding translation release factor eRF3; *su2-5* is a mutant allele of the *su2* gene encoding translation release factor eRF1 (Gagny and Silar, 1998); see text for further details about *su1* and *su2* missense mutations).
**b.** Mutated nucleotides are indicated in bold in the context of the modified codon
**c.** The *AS1-3* allele was independently isolated during the genetic selection that produced the *AS1-2* and *AS1-4* alleles and after found to be molecularly identical to *AS1-2*

The genetic screen leading to isolation of mutated alleles in the *AS1* gene is hereafter briefly redrawn. Four alleles (*AS1-1*, *AS1-2, AS1-3* and *AS1-4*) were selected as antagonizing the effect of *su1* mutations in the gene encoding translation termination factor eRF3. A fifth allele (*AS1-5*) was obtained as alleviating the growth defect associated with a *su2-5* mutation in the gene encoding translation termination factor eRF1 (Table 1 and Discussion). While *AS1-2, AS1-3* and *AS1-4* were obtained after nitrosoguanidine mutagenesis, *AS1-1* and *AS1-5* were obtained by chance. The *AS1-1* allele appeared spontaneously in a *193 su1-42* background (Table 1) and the *AS1-5* allele arose from the single *su2-5* mutant (Picard-Bennoun, 1981).

All *AS1* mutations are missense mutations (Table 1). As previously reported (Dequard-Chablat and Sellem, 1994), the *AS1-4* and *AS1-5* mutations affect the same Glycine codon substituted for an Aspartate and a Cysteine, respectively (position 139 *P. anserina* numbering; Fig. 1B). We report here that the missense *AS1-1* mutation gives rise to a Proline to Serine exchange (position 138), while the *AS1-2* and *AS1*-3 mutations are indeed identical and change a Serine to a Phenylalanine (position 145; Table 1 and Fig. 1B). Hereafter, only the *AS1-2* allele will be considered.

Alignment of the amino acid sequences of 14 representative eukaryotic r-proteins uS19 and three prokaryotic counterparts is presented Fig. 1A. *AS1* mutations modify very close residues that all fall in the C-terminal part of uS19 (Fig. 1A), precisely in the PGIGATHSSR decapeptide mapped to neighbor the A site mRNA codon (Fig. 1B) (Khairulina *et al*., 2010). Changes induced by the *AS1* mutations affect residues strictly conserved in human, *P. anserina*, and most eukaryotes. There is an exception for the phylogenetic Saccharomycotina group (exemplified here by the uS19 sequences from *S. cerevisiae*, *Candida glabrata* and *Kluyveromyces lactis*; Fig. 1A). In this group, uS19 C-terminal tail is distinguishable with a deficit of three internal residues that creates gaps in the alignment (Fig. 1A and B). Outside these gaps, the very end of all eukaryotic sequences (last eight residues) retained a high level of conservation.

### Phenotypic traits associated with the AS1 mutations define two classes of mutants

The four *AS1* mutations were studied for the phenotypes they produced on *P. anserina* organism. The present study supplements reports about the phenotypic traits associated with *AS1* mutations and especially with the *AS1-4* allele (Picard-Bennoun, 1976; Picard-Bennoun, 1981; Kieu-Ngoc and Coppin-Raynal, 1988; Belcour *et al*., 1991; Dequard-Chablat and Sellem, 1994; Contamine *et al*., 1996). Phenotypic features that characterize the development of the filamentous fungus and point out the differences between *AS1* mutant strains are summarized in Tables 2 and 3.

**Table 2.**
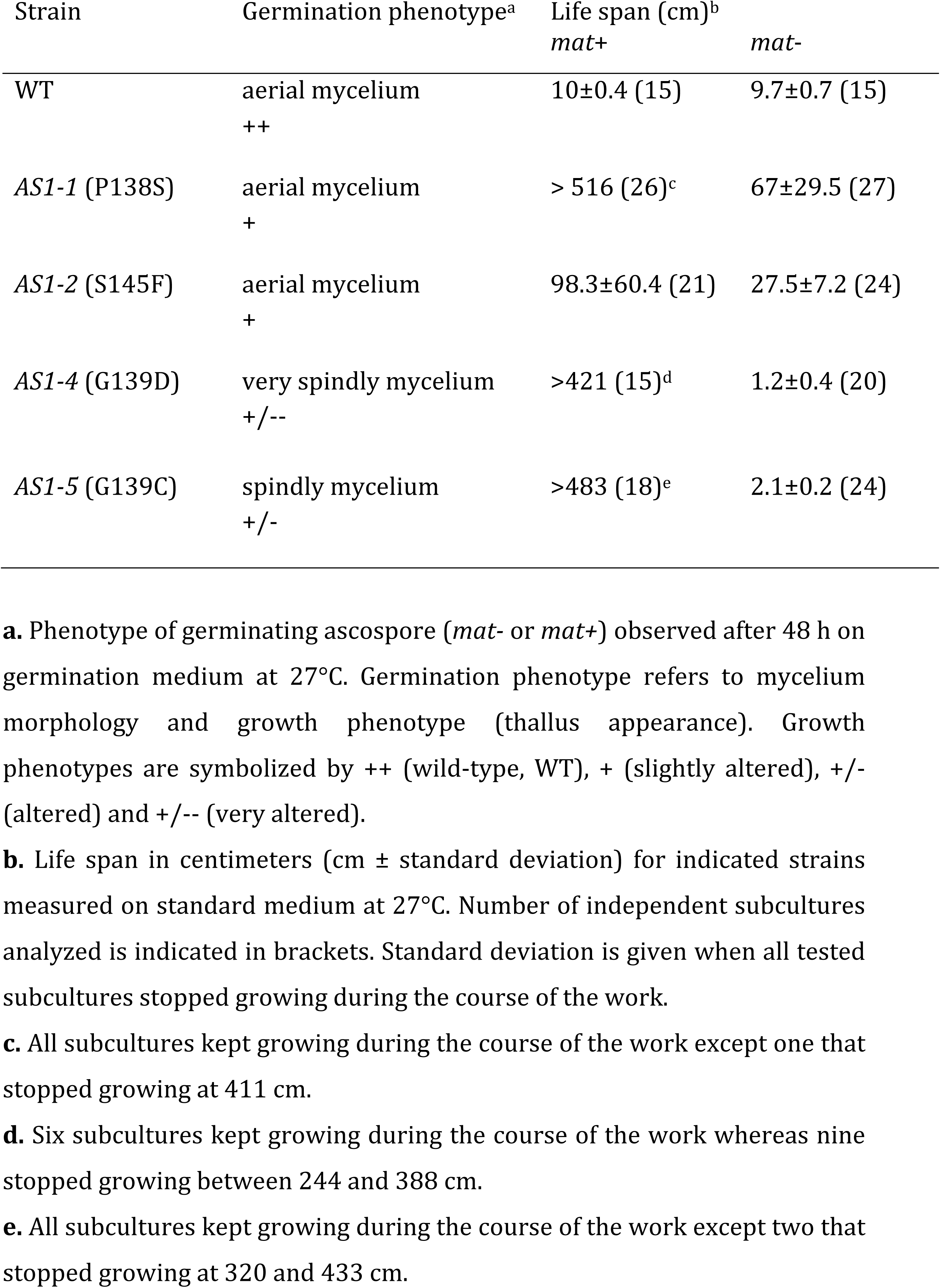
Phenotypic traits of the *AS1* mutant strains.

**Table 3.** Vegetative growth of the *AS1* mutant strains.

| Strain | Vegetative growth (cm/day)<sup>a</sup> |  |  |  |  |  |
| --- | --- | --- | --- | --- | --- | --- |
|  | 27°C |  | 11°C |  | 35°C |  |
|  | *mat*- | *mat*+ | *mat*- | *mat*+ | *mat*- | *mat*+ |
| WT | 0.72 | 0.72 | 0.15 | 0.15 | 0.84 | 0.75<sup>c</sup> |
| *AS1-1* (P138S) | 0.75 | 0.70 | 0.16 | 0.16 | 0.84 | 0.72<sup>c</sup> |
| *AS1-2* (S145F) | 0.75 | 0.72 | 0.15 | 0.16 | 0.88 | 0.73<sup>c</sup> |
| *AS1-4* (G139D) | NM<sup>b</sup> | 0.74 | 0.17 | 0.19 | 0.80 | 0.79 |
| *AS1-5* (G139C) | 0.76 | 0.74 | 0.20 | 0.20 | 0.84 | 0.80 |

**a.** Vegetative growth measured on standard medium for each indicated temperature and strain. Mycelium of *P. anserina* elongates linearly over time and growth expressed in centimeters (cm) *per* day corresponds to the slope of the linear curve obtained by following growth of at least four independent cultures *per* strain and averaging the data (see Experimental procedures for details).
**b.** Highly shortened life span of the *AS1-4 mat*- strain (see Table 2) prevents growth rate measurement at 27°C (NM= not measurable).
**c.** At 35°C, wild-type (WT), *AS1-1* and *AS1-2 mat*+ strains show a clear vegetative growth slowing down when compared to their *mat*- counterparts. Growth reduction (expressed as the (cm *per* day *mat*+/cm *per* day *mat*-) ratio in percent) is of 84%, 86% and 83%, respectively. The *AS1-5* and *AS1-4 mat*+ strains do not show such slowing down with ratios of 95% and 99%, respectively.

First, none of the *AS1* mutations affected ascospore germination that triggers the *P. anserina* life cycle (*e.g*., germination efficiency was equivalent to that of wild type). Nevertheless, the *AS1-4* and *AS1-5* germinating ascospores displayed a strong phenotype with an altered growth of the thalli and the production of spindly mycelia (Table 2). On the contrary, *AS1-1* and *AS1-2* germinating ascospores produced a wild-type aerial mycelium with only a slightly altered growth phenotype.

Life span is another phenotypic trait that characterizes the filamentous fungus *P. anserina*. At 27°C on standard growth medium (Experimental procedures), all of the *AS1* mutants showed a modified life span compared to wild type reference strains (Table 2). In the mating type *mat*+ context, all *AS1* mutants displayed a highly extended life span (10 to 50 times longer than wild type), and during the course of the work, the majority of the *mat*+ subcultures kept growing (with the exception of the *AS1-2* mutant). In the mating type *mat*-context, *AS1* mutants differ between them. On one hand, the *AS1-1* and *AS1-2* mutants still displayed an extended life span (3 to 7 times longer than wild type), 9 whereas on the other hand, *AS1-4* and *AS1-5* showed a shortened life span (5 to 8 times shorter than wild type). Life span discrepancy between the *AS1-4 mat*+ and *AS1-4 mat*-strains has been already described (Contamine *et al*., 1996) and was shown to stem from the *rmp1* gene (Contamine *et al*., 2004). That filamentous fungus specific gene is tightly linked to the mating type locus where it exists under two natural alleles: *rmp1-2* and *rmp1-1* (within the *mat*+ and *mat*-locus, respectively). The influence of *rmp1* on fungus death timing was reported for other *P. anserina* mutants but no functional explanation has been established yet (Sellem *et al*., 2005; El-Khoury and Sainsard-Chanet, 2009; Adam *et al*., 2012; see Discussion).

Finally, vegetative growth of each *AS1* mutant was examined at 27°C as well as at low (11°C) and high (35°C) temperature (Experimental procedures). Starting from 2 day germinating ascospores, five independent subcultures of each *AS1* mutant were used to calculate an average growth rate expressed in cm *per* day (Table 3). In all conditions, vegetative growth of the *AS1-1* and *AS1-2* mutants barely differed from the wild type. In particular, the wild-type, *AS1-1* and *AS1-2 mat*+ strains all exhibited an equivalent reduced growth rate at 35°C when compared to their *mat* -counterparts (Growth reduction was of 84%, 86% and 83%, respectively, see Table 3). Difference between the *mat*+ and *mat*-context is anew due to the *rmp1* gene whose *rmp1-2* allele is known to be responsible for the temperature sensitive growth of wild-type *mat*+ strain at 35°C (Contamine *et al*., 2004). Noteworthy, the *mat*+ *vs. mat*-difference in growth at 35°C was not detected for the *AS1-4* and *AS1-5* mutants (Table 3). This point will be discussed further.

Altogether, detailed phenotypic analyses revealed two classes of *AS1* mutants. First, the *AS1-1* (P138S) and *AS1-2* (S145F) mutants that differ very little from each other and 10 from the wild-type strain for their germination phenotype and vegetative growth but are however very different from wild type when examined for longevity, both mutants displaying an extended life span. In addition, life span analysis reveals a difference between the *AS1-1* and *AS1-2* mutants, the former growing longer (comparison is made for the same mating type context). Secondly, *AS1-4* (G139D) and *AS1-*5 (G139C) development is always distinguishable from the wild type and clearly different from the one of *AS1-1* and *AS1-2* mutants. The *AS1-4* and *AS1-*5 strains share similar phenotypic alterations but defects are exacerbated in *AS1-4* (especially in the *mat*-context), indicating that G139D amino acid exchange is more deleterious than G139C substitution to fungal development.

### Defects in ribosome function further pinpoint the dichotomous nature of the AS1 mutations

Considering the role of yeast uS19 in small subunit assembly (Ferreira-Cerca *et al*., 2007), we first examined whether *AS1* mutant alleles affected the production of 40S ribosomal subunit (r-subunit) in *P. anserina*. Due to the shortened life span of the *AS1-4 mat*- and *AS1-5 mat*-mutants (Table 2), experiments were all carried out in the *mat*+ context. Equivalent amounts of cell extracts from wild-type and mutant *mat*+ strains grown at 27°C on standard growth medium were fractionated on sucrose gradients in the absence of Mg2^+^, to dissociate ribosomes and polysomes into free 40S and 60S r-subunits (Experimental procedures). *AS1* mutations led to either modest (*AS1-1* and *AS1-2*) or clear (*AS1-4* and *AS1-5*) reduction in the amounts of free 40S r-subunits relative to 60S r-subunits (Fig. 2 bottom panels). Decrease in 40S r-subunits was further estimated by calculating 40S-to-60S ratios for at least three independent sucrose gradients using two independent extractions for each strain (Fig. 2 histograms). In agreement with the phenotype traits associated with *AS1* mutants, *AS1-4* showed the 11 strongest deficit in 40S r-subunits with a 40-to-60S ratio of 41% in comparison to 69% for the wild-type strain. *AS1-1* and *AS1-2* displayed equivalently slightly reduced 40S-to-60S ratios (60%) when in the *AS1-5* mutant the mean value of 40S-to-60S ratio is 49%. We next examined the effect of 40S/60S subunit imbalance on polysome profiles whose analysis is a sensitive method for detecting defects in translation. Polysome profiles were compared for equivalent amounts of extracts from wild type and mutant *mat*+ strains grown also at 27°C on standard growth medium (Fig. 3A) (Experimental procedures). Overall, *AS1-1* and *AS1-2* strains gave rise to gradient profiles similar to that of wild type with equivalent peaks of free 40S, free 60S, monosomes (80S) and polysomes (the grid pattern in Fig. 3A helps the comparison). Polysomes content of the *AS1-5* mutant was also comparable to that of wild type whereas much less translating ribosomes (polysomes) were detected for the *AS1-4* strain. Consistent with the reduced amount of 40S produced in *AS1-4* (Fig. 2), no peak of free 40S r-subunit was observable on *AS1-4* gradient profiles along with a concomitant large excess of free 60S r-subunit (Fig 3A). Similarly, but to a lesser extent, the *AS1-5* gradient profiles hardly showed a peak of free 40S while the peak of free 60S increased in comparison to wild type.

**Fig. 2.**
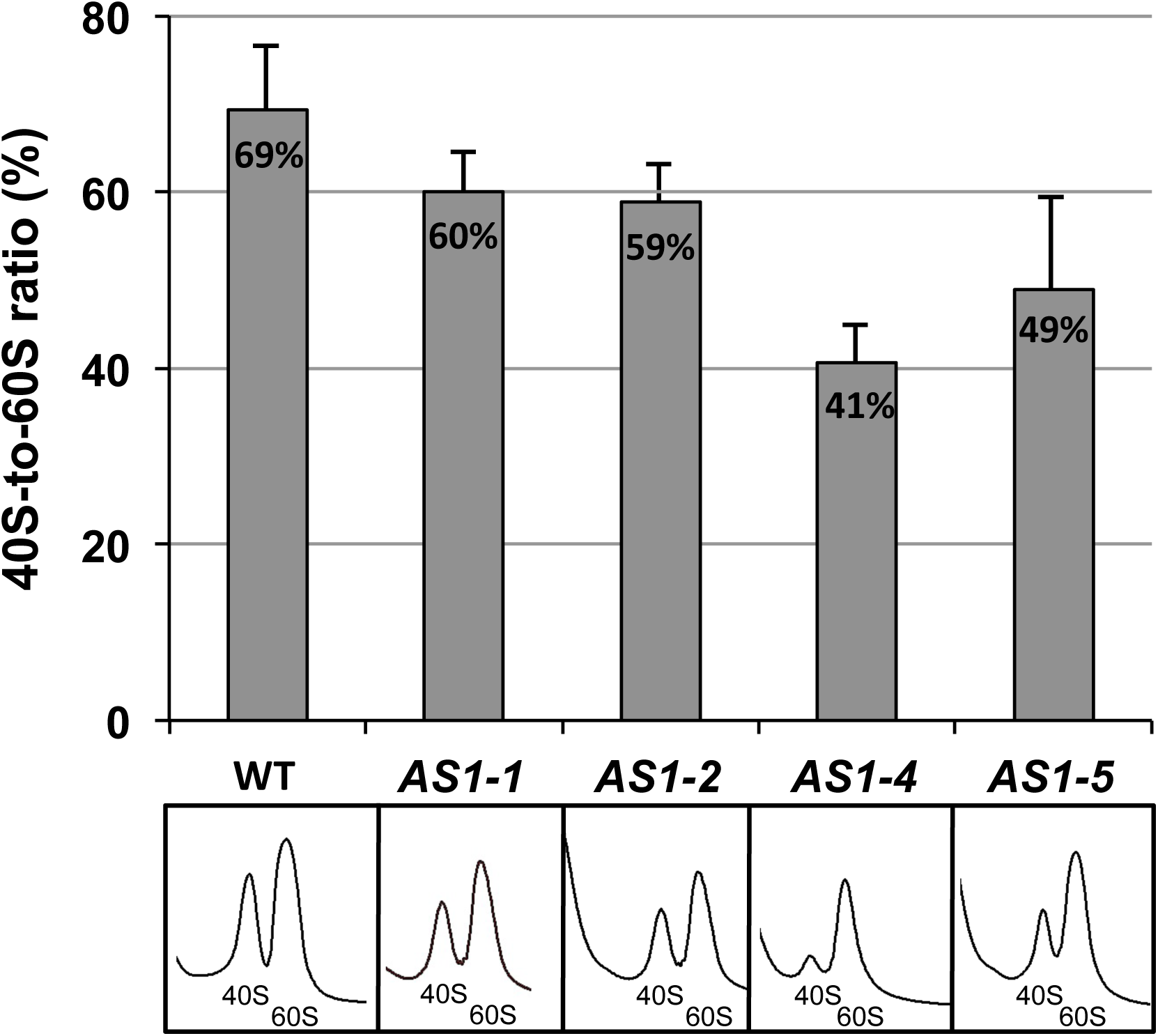
Production of 40S subunits is modestly to strongly reduced in *P. anserina AS1* mutants. Equivalent amounts of cell extracts (10 A_260nm_ units) obtained from wild-type and mutant *mat*+ strains grown at 27°C on standard growth medium were fractionated on 7%-50% sucrose gradients. Representative profiles of gradients analyzed by continuous monitoring at A_254nm_ are shown (bottom). At least three independent sucrose gradients and two independent extractions for each strain were used to quantify the 40-to-60S ratios (Top). Ratios were determined by measuring the area under the 40S peak related to the area under the 60S peak. For each strain, the 40S-to-60S ratios are expressed in percentage and error bars represent the standard deviation of the mean. Significance of the differences between mutants and wild type was calculated using a parametric *t*-test (XLSTAT) *P*-values of 0.0004 (very highly significant) and 0.027 (significant) were obtained for *AS1-4* and *AS1-5*, respectively while for *AS1-1* and *AS1-2* less confidence difference was measured (*P* = 0.104 and *P* = 0.07, respectively).

**Fig. 3.**
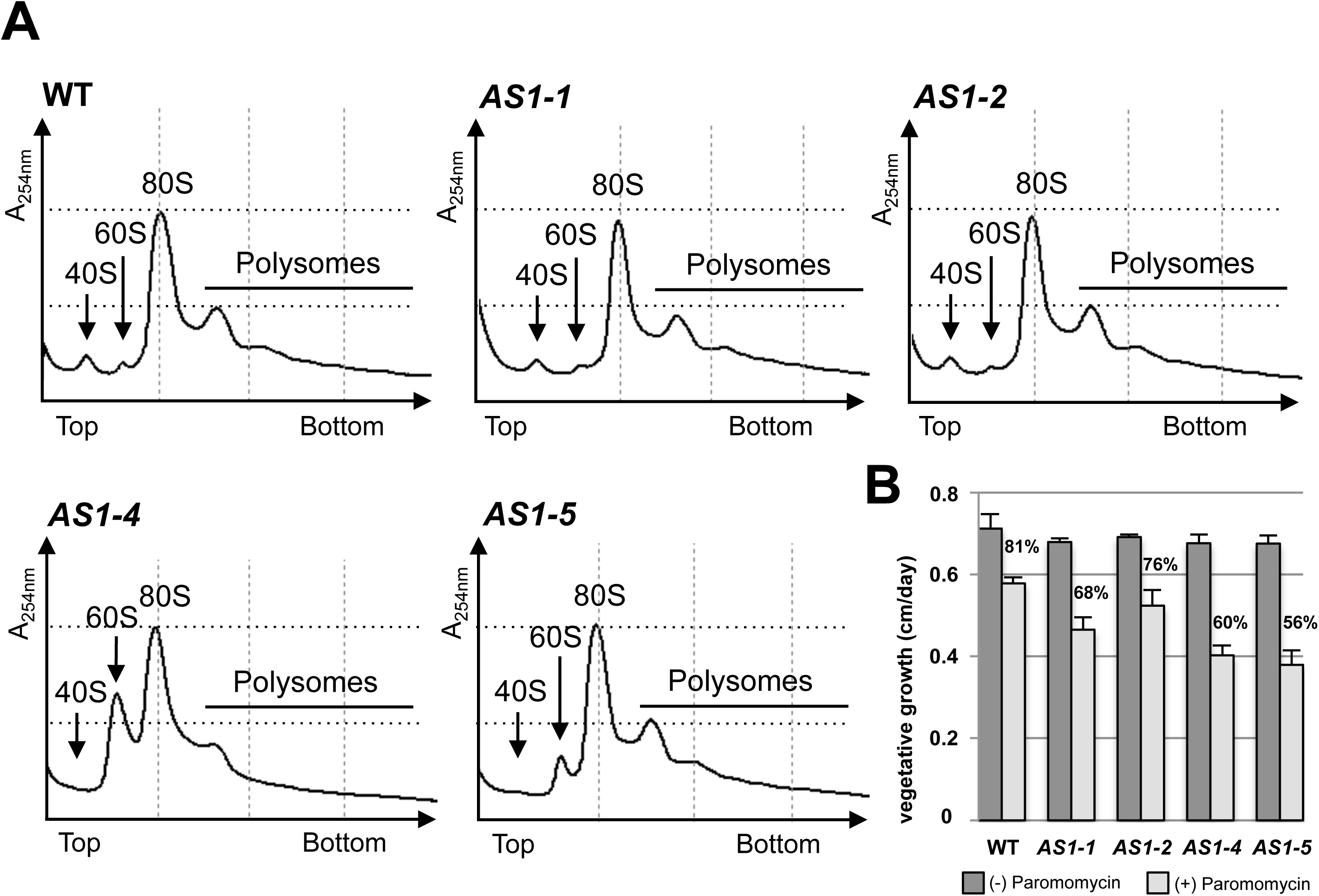
Polysome profiles and paromomycin sensitivity reveal two classes of *P. anserina AS1* mutants. A. Representative polysome profiles obtained for wild-type and mutant *mat*+ strains grown at 27°C on standard growth medium. Equivalent amount (10 A_260nm_ units) of cycloheximide-treated mycelium extracts were fractionated on 7%-50% sucrose gradients containing Mg^2+^. Gradients were analyzed by continuous monitoring at A_254nm_ from top to the bottom. Positions of free 40S, free 60S, 80S monosomes and polysomes (translating ribosomes) are indicated. Arrows point to the 40S and 60S peaks or to their expected positions in the gradient. Dotted grid pattern guides profile comparison and highlights *AS1-5* and *AS1-4* alterations. B. Paromomycin effect on vegetative growth of wild-type and *AS1* mutant *mat*+ strains. Four subcultures of each strain were grown at 27°C on standard growth medium containing (+) or not (-) 500 μg ml^-1^ of drug. Mycelium growth was followed during nine days. For each strain and condition, vegetative growth was measured in cm/day (Experimental procedures) and data are expressed as mean values ± standard deviation. Average percentage of residual growth in the presence of drug (+) relative to growth in its absence (-) is indicated above the histogram for each strain. Significance of the differences between mutants and wild type was calculated using a parametric *t*-test (*P* = 0.06 (*AS1-2*), *P* ≤ 0.003 (*AS1-1; AS1-4; AS1-5*)).

Defects in translation can be also detected by testing sensitivity to paromomycin, a translation error-inducing aminoglycoside antibiotic. Growth rate of the wild type and mutant *mat*+ strains were compared during nine days of vegetative growth at 27°C on standard medium with or without 500 μg ml^-1^ paromomycin (see Experimental procedures). For each strain, growth rate of the mycelium on drug-containing medium was expressed as the percentage of its growth rate on drug-free medium (Fig. 3B). In the presence of paromomycin, growth rate of the wild-type strain was 81% compared to plain media. Consistent with previous independent report (Kieu-Ngoc and Coppin-Raynal, 1988), the *AS1-2* mutant displayed a paromomycin-sensitivity barely different 12 from the wild type (76% of residual growth) while the *AS1-1* mutant exhibited a slight hypersensitivity (68 % of residual growth). In comparison, the *AS1-4* (this work and Contamine *et al*., 1996) and *AS1-5* (this work) mutants were both found hypersensitive to paromomycin, displaying growth rates almost half reduced.

Just like ribosome defects, comparison of paromomycin sensitivity differentiates *P. anserina AS1* mutants into two classes: *AS1-1* and *AS1-2* on one a hand and *AS1-4* and *AS1-5* on the other hand. The *AS1-4* (G139D) mutant displays the strongest fungal development defect and the strongest ribosomal alterations. In accordance with phenotypic analyses, the *AS1.1* (P138S) and *AS1.2* (S145F) mutants show light alteration in 40S production with no detectable consequences on polysomes while *AS1-5* (G139C) exhibits intermediary ribosome defects.

### ScAS1-4 (G139D) mimetic mutation induces growth and ribosomal defects in the yeast S. cerevisiae

To further evaluate the translation defects associated with mutations affecting the C-terminal tail of uS19 r-protein, the four *P. anserina AS1* mutants were mimicked in yeast *S. cerevisiae.* As mentioned before, primary sequence of *S. cerevisiae* uS19 C-terminal tail is quite different from that of *P. anserina* especially in the decapeptide motif 138-PGIGATHSSR-147 (Fig. 1A and B). Because the difference between yeast and *P. anserina* sequences prevents copycatting of the *AS1* mutations in the natural *S. cerevisiae RPS15* gene, *AS1* mutations were introduced in a *RPS15* chimeric gene designed to code for an yeast r-protein uS19, whose last 25 amino acids have been exchanged for the last 28 amino acids of the *P. anserina* polypeptide (Fig. 1B; see Experimental procedures for details). Each chimeric *RPS15* gene was put under the control of the endogenous yeast *RPS15* promoter and expressed from a low copy vector that was introduced in an *rps15* deleted yeast strain using plasmid shuffling. Resulting *S. cerevisiae* strains harboring the *AS1-1*, *AS1-2*, *AS1-4* or *AS1-5* mutations were named Sc*AS1-1*, Sc*AS1-2*, Sc*AS1-4* and Sc*AS1-5*, respectively. In addition, the Sc*AS1* strain refers to the same *rps15* deleted strain that expresses the wild-type version of the chimeric uS19 r-protein, while ScWT is the *rps15* deleted strain expressing the natural yeast *RPS15* gene from the same low copy vector.

The resulting six yeast strains were compared for their vegetative growth on glucose rich medium (Fig. 4A). At all tested temperatures, growth of the Sc*AS1* strain was equivalent to that of the ScWT reference strain indicating that the chimeric uS19 protein can functionally replace the endogenous yeast small r-protein and that albeit divergent in size and sequence, the C-terminal tail of *P. anserina* uS19 can substitute for that of the *S. cerevisiae* protein. Of note, in the course of chimera constructions (see Experimental procedures), we found that the deletion of the very last 15 aas of the chimeric uS19 r-protein was lethal to yeast, which further highlights the important role played by the C-terminal tail of this eukaryotic r-protein.

**Fig. 4.**
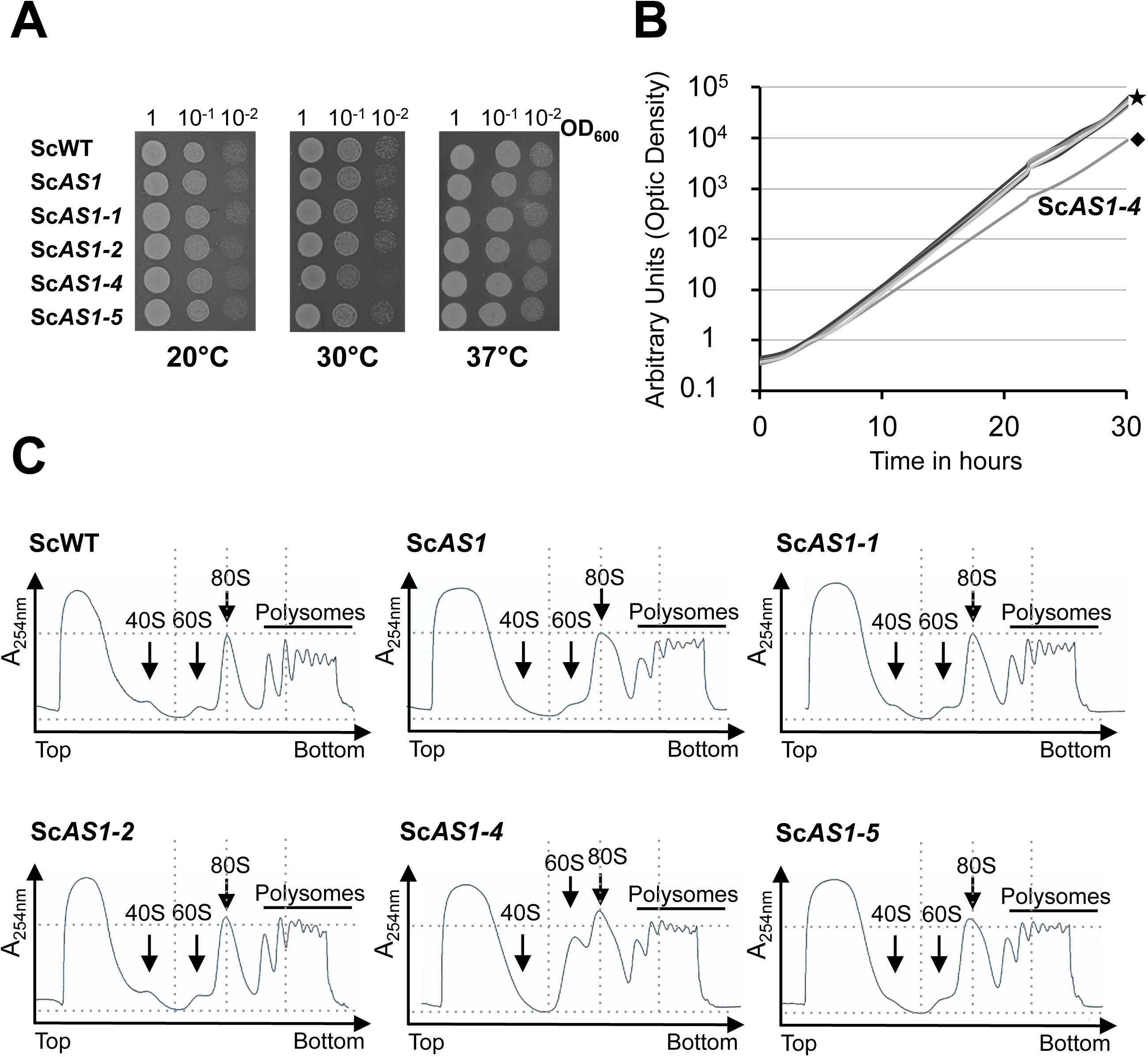
Growth and polysome profiles of the *S. cerevisiae* Sc*AS1* mimetic mutants distinguish the Sc*AS1-4* strain. A. Growth of equivalent 10-fold serial dilutions (indicating OD_600_) of the indicated strains after 2 days (20°C) or 1 day (30°C, 37°C) on glucose rich medium (see Fig S1). ScWT, reference strain expressing the wild-type *S. cerevisiae RPS15* gene; Sc*AS1*, strain expressing a wild-type *RPS15* chimeric gene in which last 75 nucleotides have been replaced by the last 84 nucleotides of *P. anserina AS1* gene (Fig. 1B); Sc*AS1-1* to Sc*AS1-4*, yeast strains mimicking the *AS1-1* to *AS1-4 P. anserina* mutations, respectively. B. Growth curves of the same strains grown at 30°C in glucose rich medium. Apart from Sc*AS1-4* (diamond), Sc*AS1* and other mimetic strains show growth curves that merge with that of the ScWT reference strain (star). C. Representative polysome profiles obtained for the indicated strains grown at 30°C in glucose rich medium. Equivalent amount of cycloheximide-treated yeast extracts were fractionated on 7 to 50% sucrose gradients. Gradients were analyzed by continuous monitoring at A_254nm_ from top to the bottom. The positions of the different ribosomal species are indicated. Apart from the Sc*AS1-4* strain, which displays no 40S peak and an excess of free 60S (as its *P. anserina* counterpart, Fig. 3A), polysome profiles of Sc*AS1* and other mimetic strains are close to that of the ScWT reference strain.

All mimetic yeast mutants supported wild-type growth on solid glucose rich medium at all tested temperatures (Fig. 4A). When tested for growth at 30°C in liquid glucose rich medium, the Sc*AS1-4* mutant stood out from the others and displayed a clear slowing down of growth after 30 hours of culture (Fig. 4B). On solid medium at 30°C, an altered growth of the Sc*AS1-4* strain could be also observed one day after the incubation of serial dilutions (see Fig. 4A) but this difference was no more detectable after two days of incubation (Fig. S1). We never observed such a growth delay at low (20°C) or high (37°C) temperature for the Sc*AS1-4* strain or the other mutant strains (Fig. 4A and Fig. S1).

We next examined the polysome profiles for equivalent amounts of yeast extracts from all six strains grown at 30°C in glucose rich medium (Fig. 4C). Apart from the Sc*AS1-4* mutation which gave rise to an excess of free 60S subunit and no detectable peak of free 40S subunit as observed for the *P. anserina AS1-4* strain, the polysome profiles of the *ScAS1* strain and of the Sc*AS1-1*, Sc*AS1-2* and Sc*AS1-5* mimetic mutants did not significantly differ from that of the ScWT reference strain (Fig. 4C).

Compared to the complex phenotypes associated with the presence of the *AS1* mutations in the multicellular filamentous fungus *P. anserina*, the mimetic Sc*AS1* mutations gave rise to very modest effects in the unicellular yeast *S. cerevisiae* (within the limits of the phenotypes tested), suggesting that ribosomal proteins are not equally important for fungi development (Romani *et al*., 2012).

### Depending on uS19 tail alterations, stop codon recognition is differently affected in yeast mimetic mutants

Considering that *P. anserina AS1* mutations were selected as translational control mutations, we next investigated whether stop codon recognition could be affected in yeast mimetic mutants, despite the absence of strong growth phenotype. To monitor potential stop codon readthrough, we employed a widely used dicistronic *LacZ-luc* dual-gene reporter system where one stop codon (UAG, UAA, or UGA) is inserted between the *E. coli LacZ* and firely luciferase (*luc*) open reading frames so that firely luciferase can only be produced as a result of nonsense suppression (Fig. 5A). Plasmids bearing the dual-gene reporter system were introduced in the reference and mimetic mutant strains and readthrough assays were performed using crude extract of exponentially growing cultures in glucose medium at 30°C (Experimental procedures).

**Fig. 5.**
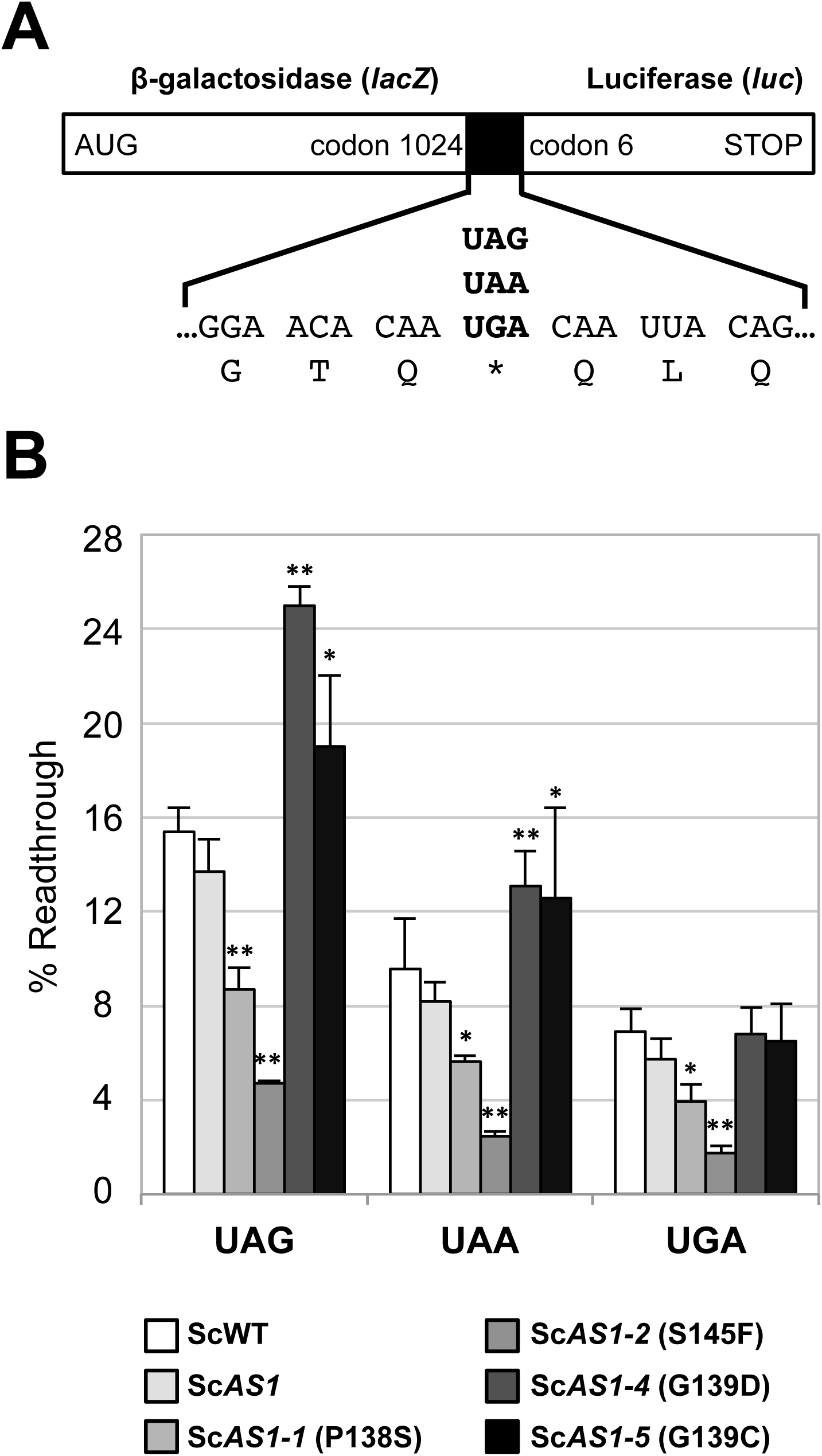
Termination codon readthrough is differently altered in the *S. cerevisiae* Sc*AS1* mimetic mutants. A. Schematic representation of the β-galactosidase (*lacZ*)-luciferase (*luc*) dual-reporter system used. Positioning of the nonsense-containing region within the *lacZ*-*luc* dual-gene is indicated by the black box and bordering sense codons of *lacZ* and *luc* open reading frames. Only a section of the nonsense-containing sequence (21 nucleotides among 51; Experimental procedures) is given. B. Quantification of nonsense codon readthrough in each indicated strain grown at 30°C in glucose selective medium (bottom). Strains were transformed with control, UAG, UAA and UGA *LacZ-luc* reporter plasmids and the percent readthrough in each strain was expressed as the ratio of luciferase activity (measured for the indicated nonsense codon) to the one measured for an in frame *LacZ-luc* construct (control), after β-galactosidase normalization (Experimental procedures). Bars represent mean ± standard deviation and means were determined for at least five independent measurements. The significance of differences in signals between each mimetic mutant and the Sc*AS1* strain is indicated: \**P* < 0.02, \*\**P* < 0.002 (parametric *t*-test).

The stop codon containing-sequence of the *LacZ-luc* dual-gene reporter system (Fig. 5A) was previously shown to promote high level of readthrough in yeast (Stahl *et al*., 1995; Bidou *et al*., 2000). Accordingly, we measured a readthrough level of 7 % (UGA) to 15% (UAG) in the ScWT reference strain expressing the natural yeast *RPS15* gene (Fig. 5B; see Experimental procedures). In the Sc*AS1* strain, whatever the stop codon the readthrough level was close to that of the ScWT strain, indicating that wild-type chimeric uS19 r-protein did not modify stop codon recognition in yeast. On the contrary, all four mimetic mutants exhibited altered stop codon recognition (Fig. 5B). When compared to Sc*AS1*, the Sc*AS1-1* and *ScAS1-2* mutant strains displayed hyperaccurate recognition of the three stop codons and, whatever the stop codon, an approximately 1.5 and 3-fold enhancement was found in the Sc*AS1-1* and Sc*AS1-2* strains, respectively (Fig. 5B). On the opposite, the Sc*AS1-4* and Sc*AS1-5* mutant strains both exhibited higher readthrough at the UAG and UAA codons whereas no significant change was detected in the recognition of the UGA codon. In the Sc*AS1-4* strain, readthrough level increased from 1.6- (for the UAA codon) to 1.8-fold (for the UAG codon) when compared to that in the Sc*AS1* strain (Fig. 5B). In the Sc*AS1-5* strain, UAG and UAA readthroughs are increased approximately 1.4- and 1.5-fold, respectively.

Our readthrough assays strongly support that the C-terminal tail of uS19 contributes *in vivo* to the recognition of the different stop codons. These results moreover revealed that depending on residues modified in the decapeptide PGIGATHSSR, translation termination ambiguity could be either expanded or on the contrary restricted, which further emphasizes the key role played by this small set of amino acids within the C-terminal extension of eukaryotic uS19 r-protein.

## Discussion

S15/uS19 is an essential r-protein and one of the fifteen universally conserved ribosomal proteins of the eukaryotic small ribosomal subunit. *In vitro* cross-linking studies using mammalian 40S and 80S complexes have identified uS19 as the r-protein that intimately contacts -*via* its C-terminal tail- the mRNA at the A site codon during translation, making uS19 a key component of the eukaryotic decoding site (Graifer *et al*., 2004; Bulygin *et al*., 2005; Molotkov *et al*., 2006; Pisarev *et al*., 2006; Pisarev *et al*., 2008). Further investigations into the role of uS19 during the translation process however suffered from the absence of functional analyses and the non-resolution of the structure of the C-terminal tail.

In this study, we looked back on genetic screens that four decades ago had identified the uS19 encoding-gene *AS1* as an actor of the translation fidelity in the filamentous fungus *P. anserina* (Picard, 1973; Picard-Bennoun, 1976; Picard-Bennoun, 1981). All *AS1* mutations isolated then are now identified and described hereinbefore as altering conserved residues of uS19 C-terminal tail and more precisely three residues of the eukaryotic specific peptide 138-**PG**IGATH**S**SR-147 (modified residues in bold, *P. anserina* numbering), which was shown *in vitro* to be the contact site of the mRNA during the elongation and termination steps of translation (Khairulina *et al*., 2010).

To *in vivo* investigate the translational role of the PGIGATHSSR decapeptide, we have considered a widely used method based on a dicistronic *LacZ-luc* dual-gene reporter system and measure translational error rates in yeast *S. cerevisiae*. All *P. anserina AS1* mutations were recreated within the yeast counterpart uS19 encoding-gene and stop codon readthrough monitoring revealed that all resulting substitutions (P138S, G139D, G139C and S145F) significantly alter the termination process in yeast. Regardless of the nature of the stop codon, P138S and S145F changes both significantly reduce readthrough level (*e.g.,* increase translational accuracy) whereas substitutions of the G139 residue display an opposite effect and slacken stop codon recognition at UAG and UAA (*e.g.,* increase translational error). No significant modification of UGA recognition was detectable for G139D or G139C changes suggesting that this glycine residue plays a role in discriminating between the presence of a Guanine or an Adenine at the second position of the stop codon (*i.e*., the second nucleotide within the A site). Consistently, cross-linking studies have shown that in mammalian ribosome complexes the C-terminal tail of uS19 mainly contacts the first and second nucleotide of the A site (Khairulina *et al*., 2010; Graifer and Karpova, 2012; Sharifulin *et al*., 2015). In the absence of a solved structure for eukaryotic uS19 C-terminal tail, a modeling has been proposed that placed the last four residues of the decapeptide 138–PGIGATHSSR-147 in close contact with the first and second position of the A site (Khairulina *et al*., 2010). Even though translational effect associated with the S145P change suggests an *in vivo* proximity of this residue with the A site, the proposed positioning of the decapeptide in this model does not place G139 nor P138 close to it whereas our *in vivo* results clearly demonstrate an important, while opposite, implication of these two residues in stop codon recognition. A short distance (intra-peptide) influence, either structural or functional, cannot be excluded.

Considering the genetic screen initially designed to isolate *AS* (<u>a</u>nti-<u>s</u>uppressor) mutations in *P. anserina* we could have expected all *AS1* mutations to increase translational accuracy. However in yeast, only two *AS1* mutations (Sc*AS1-1*(P138S) and Sc*AS1-2*(S145F)) gave rise to an hyperaccurate recognition of the stop codons whereas alterations of the G139 residue displayed clearly an opposite effect. Such discrepancy could result from the presence in *P. anserina* of *su* (<u>su</u>ppressor) mutations at the time of *AS* genetic isolation (Picard-Bennoun, 1976; Belcour *et al*., 1991; Dequard-Chablat and Sellem, 1994). Alternatively, we cannot exclude that in yeast the mimicking mutations had different effects during the termination process all the more because the PGIGATHSSR decapeptide is not conserved in the *S. cerevisiae* uS19 r-protein (Fig. 1). Nevertheless, with the C-terminal tail from *P. anserina*, *S. cerevisiae* uS19 chimeric r-protein turned out to be functional in yeast and did not alter stop codon recognition. Noteworthy, phenotypic traits associated with *P. anserina AS1* mutations suggest that cleared off the presence of the *su* mutations, the *AS1-4* and *AS1-5* mutations might promote *in vivo* readthrough on UAG codon as observed in yeast. To get to this conclusion, we paid attention to the very low or absence of temperature sensitivity of *P. anserina AS1-4* and *AS1-5 mat*+ strains compared to wild-type, *AS1-1* and *AS1-2 mat*+ strains (Table 3). As previously mentioned, the *rmp1-2* allele of the mating-type-linked *rmp1* gene is responsible for the temperature sensitive growth of *mat*+ strains compared to *mat*-strains (Contamine *et al*., 2004). Compared to the *rmp1-1* allele (present in *mat-* strain, dominant over *rmp1-2* and considered as fully functional), the sequence of the *rmp1-2* allele (present in *mat*+ strain) contains two nucleotide modifications: a functionally neutral missense mutation and a nonsense mutation (UAG) that truncates the very last 19 amino acids of RMP1 polypeptide (Contamine *et al*., 2004). By promoting readthrough on UAG, the *AS1-4* (G139D) and *AS1-5* (G139C) mutations might restore synthesis of some full length RMP1 polypeptide thereby accounting for the relief of the temperature sensitivity observed for both *P. anserina AS1-4* and *AS1-5 mat*+ strains (Table 3). While indirect that observation nevertheless suggests that in *P. anserina*, substitutions of the G139 residue could increase translational error as shown in *S. cerevisiae*.

Importance of the C-terminal tail of uS19 r-protein to stop codon recognition is further supported by the fact that *P. anserina AS1* mutations were isolated as interacting with mutations in the genes coding for the release factors eRF1 and eRF3, whose interdependent interaction mediates eukaryotic translation termination. eRF3 protein is a GTPase that forms a ternary complex with GTP and eRF1, delivering eRF1 to the A site. In the A site, eRF1 is responsible for recognition of all three stop codons whereas eRF3’s GTPase activity enhances polypeptide release (Hellen, 2018, for review). As described hereinbefore, the *AS1-1* (P138S), *AS1-2* (S145F) and *AS1-4* (G139D) mutations display the property to diminish the non-sense suppressor mutation efficiency of a *P. anserina* eRF3 mutant that contains a unique S380N substitution in the GTPase domain of the protein (Picard, 1973; Picard-Bennoun, 1976; Belcour *et al*., 1991; This work, see Table 1). Serine 380 residue (*P. anserina* numbering; Serine 319 in *Schizosaccharomyces pombe*; Threonine 293 in human eRF3A) is located between the switch I and switch II elements that are present in all GTPases and essential for binding and hydrolysis of GTP (Kong *et al*., 2004; Cheng *et al*., 2009). In translation pre-termination complex, eRF1 was shown to contact both the switch I and switch II regions of eRF3 and that interaction was proposed to tighten the contact between eRF1, eRF3 and the small ribosomal subunit during the decoding of the stop codon (Preis *et al*., 2014). Taking all together, we could assume that the small r-protein uS19 contributes to that interaction through its C-terminal tail. While *P. anserina* S380N eRF3 mutant is associated with loose stop codon recognition, presence of P138S, G139D or S145F substitutions in uS19 can antagonize the effect of S380N change. Thus, as an r-protein neighboring the A decoding site (Bulygin *et al*., 2002; Bulygin *et al*., 2005; Pisarev *et al*., 2006; Khaĭrulina *et al*., 2008; Pisarev *et al*., 2008; Khairulina *et al*., 2010), uS19 could modulate eRF3-eRF1 interaction in the pre-termination complex, thereby participating in stop codon recognition.

The fourth *P. anserina* uS19 mutant (*AS1-5*, G139C) was isolated as alleviating the growth defect of an eRF1 mutant that has a F407S substitution in the C-terminal domain of the protein (Dequard-Chablat and Sellem, 1994; M. Picard, unpubl. work; This work). eRF1 C-terminal domain is mainly responsible for the eRF1-eRF3 interaction (Hellen, 2018) and the highly conserved Phenylalanine 407 residue (*P. anserina* numbering; residue 405 in *S. pombe*; 406 in human eRF1) belongs to a hydrophobic patch that was shown to interact with hydrophobic residues of the C-terminal domain of eRF3 in the crystal structures of human and *S. pombe* eRF1-eRF3 complexes (Cheng *et al*., 2009). Interestingly, it has been shown that substitution of F405 to alanine in *S. pombe* eRF1 affected cell growth and strongly reduced eRF1 binding to eRF3 (Cheng *et al*., 2009). How G139C uS19 mutant alleviates growth defect of an F407S eRF1 mutant in the filamentous fungus *P. anserina* remains an opened question but it could be proposed that in the pre-termination complex, the G139C change re-establishes a functional eRF1-eRF3 interaction that has been weakened by the F407S change. Although this hypothesis deserves additional experiments, it anew indicates that uS19 C-terminal tail is an important component of the eukaryotic decoding site.

In the end, one might argue that translation accuracy is an exquisite balance between numerous, potentially competing, chemical interactions (e.g. cognate, near-cognate, non cognate codon-anticodons interactions, competing with eRFs binding to the A site), and mimicking mutations outside their natural host might be a difficult way to decipher a precise role for an individual residue. Nevertheless, we clearly identified a translation/termination defect in yeast mutants, showing a conserved role for uS19 in controlling A site interactions, to notably the nucleotide level, as G139 discriminates the presence of and Adenine or a Guanine in the second position of a stop codon. To the best of our knowledge this is an un-described fine tuning role for a ribosomal protein.

Finally, human uS19-encoding gene was recently reported as a putative cancer driver whose low frequency mutations are associated with poor outcome for patients with relapsed chronic lymphocytic leukemia (CLL; Ljungström *et al*., 2016; Bretones *et al*., 2018). Some recurrent mutations cluster at the C-terminal tail of the human protein and two of them induce the exact same changes as *P. anserina AS1-1* and *AS1-2* mutations (P131S and S138F; *H. sapiens* numbering). What is the causative effect of ribosomal uS19 mutations on CLL pathobiology and how uS19 mutations provoke pleiotropic effects on growth and development of the filamentous fungus *P. anserina* remain largely unknown but alteration of cell proteomes could probably be a part of (Bretones *et al*., 2018).

## Experimental Procedures

### *P. anserina* strains and culture conditions

All *Podospora anserina* strains used in this study derived from the S (Uppercase S) wild-type strain (Rizet, 1952; Picard-Bennoun, 1976) The selection modes of the *AS1* mutants were originally described in (Picard-Bennoun, 1976) or unpublished (for *AS1-5*). Since selection modes are important to the purpose of the present work they were recalled and detailed in the text (see Results and Table 1). Selection of *su* mutants was originally described in (Picard, 1973) and the *su1* and *su2* genes further identified as encoding the eRF3 and eRF1 translation termination factors, respectively (Gagny and Silar, 1998). Mutant *193* was described initially in (Picard, 1971) and molecularly analysed in (Coppin and Silar, 2007).

Standard culture conditions, media and genetic methods can be accessed at http://podospora.i2bc.paris-saclay.fr. The germination phenotype of germinating ascopores (Table 2) was observed on germination (G) medium after two days at 27°C. Vegetative growth and life span measurements were carried out on M2 standard medium containing mainly dextrin as a carbon source. To do so, small pieces of mycelium developed onto G medium were transferred to M2. For growth rate measurements (Table 3), depending on the temperature of growth, four (11°C) to seven (27°C, 35°C) measurements were done over time following the growth of 4-to-6 independent cultures for each strain during seven (27°C, 35°C) to fourteen days (11°C). For each strain, a linear curve was then established using averaged daily-measurements (27°C, 35°C) or averaged measurements equally distributed over 14 days (11°C). The slope of each curve (in cm *per* day) represents the vegetative growth of each strain in the indicated culture conditions.

Paromomycin sensitivity was evaluated at 27°C by following over nine days the growth of four subcultures of wild-type and mutant *mat*+ strains on M2 medium supplemented (or not) with 500 μg ml^-1^ of paromomycin (Sigma). For each strain, each subculture and condition, a linear curve was established and the slope of each curve displays vegetative growth in cm/day. The average and standard deviation of each set of data were plotted onto Fig. 3B.

### Molecular analysis of the su1-1, su1-42 and su2-5 mutants

The genomic DNA from *su1-1*, *su1-42* and *su2-5* mutants was extracted from cultures grown on cellophane disks placed on M2 medium (27°C, two days). After mycelium crushing in a FastPrep-24 machine, extractions used the ZR Fungal/Bacterial DNA miniprep Kit (Proteigene). PCR amplification with extracted DNA was carried out with primers su1-5’ and su1-3’rev for the *su1* alleles and primers su2-5’ and su2-3’rev for the *su2-5* allele (for all primers used in this study, see Table S1). PCR fragments were purified by PEG precipitation and directly sequenced on both strands (Beckman Coulter Genomics France) using oligonucleotides su1-5’, su1-3’rev, su1-int, su1-int-rev, su2-5’, su2-3’rev, su2-int and su2-int-rev. DNA extraction, PCR amplification and DNA sequencing were carried out from two independent subcultures for each mutant strain. The *su1-1* and *su1-42* alleles were found to be molecularly identical and carry one nucleotide substitution in codon 380 (*P. anserina* numbering for eRF3) that induces a Serine (A**G**T) to Asparagine change (A**A**T). The *su2-5* allele carries one nucleotide substitution in codon 407 (*P. anserina* numbering for eRF1) that causes a Phenylalanine (T**T**T) to Serine change (T**C**T).

### *S. cerevisiae* strains, ScAS1 mimetic mutants and growth conditions

*Saccharomyces cerevisiae* strain ME14-a9 (*MATa his3Δ1 leu2Δ0 lys2Δ0 ura3Δ0 MET15 rps15::kanMX4* [pFL38-*GAL::RPS15*]) (see Léger-Silvestre *et al*., 2004) was used as the recipient strain for mimicking the *P. anserina AS1* mutants. ME14-a9 is derived from Euroscarf strain Y21731 (BY4743 background). In ME14-a9, the chromosomal copy of *RPS15* is inactivated by a *kanMX4* cassette and strain viability is supported by a plasmid-borne copy of *RPS15*, which is under the control of the conditional *GAL1/10-CYC1* promoter (pFL38-*GAL::RPS15, URA3, CEN*).

Strains were grown on either solid or liquid form of glucose rich or glucose selective medium at indicated temperatures. Where required, 5-fluoroacetic acid (5-FOA) and G418 were used at final concentrations of 200 μg ml^-1^ and 100 μg ml^-1^, respectively. Plasmids pFL36 (*LEU2*) and pFL38 (*URA3*) used in this work are two low copy (*CEN*) vectors (Bonneaud *et al*., 1991). Sc*AS1* chimeric genes were initially constructed in plasmid pFL38-*RPS15wt* (Bellemer *et al*., 2010), which contains the promoter and coding sequence (CDS) of wild-type *RPS15* gene followed by a *PGK1* terminator sequence. First, we took benefit of the presence of two nearby *Eco*RI restriction sites to manipulate the 5’ end of *RPS15* CDS in order to construct *S. cerevisiae*/*P. anserina* chimeric alleles. Yeast *RPS15* CDS contains one natural *Eco*RI site at position 352-356 encompassing codons 118 and 119 (*S. cerevisiae* numbering; codon 125 and 126, *P. anserina* numbering; see Fig. 1B). The second *Eco*RI site used is located in pFL38-*RPS15wt* sequence, 12 bp downstream the *RPS15* stop codon. An internal deletion of the *Eco*RI-*Eco*RI fragment of pFL38-*RPS15wt* generated plasmid yEPU-*RPS15*-ΔC. A wild-type *S. cerevisiae*/*P. anserina* chimeric gene (Sc*AS1*) was first constructed by replacing the *Eco*RI-*Eco*RI fragment of pFL38-*RPS15wt* by an *Eco*RI-*Eco*RI fragment amplified from pLF61-7D using primers oAS1Eco-Fwd and oAS1-Eco-Rev. Plasmid pFL61-7D originates from a *P*. *anserina* cDNA library constructed in the pFL61 vector ((Espagne *et al*., 2008)). Resulting plasmid yEPU-*RPS15*-*AS1* expresses the wild-type (WT) version of chimeric uS19 r-protein in which the last 25 amino acids (aas) of *S. cerevisiae* polypeptide have been switched for the last 28 aas of *P. anserina* r-protein (Fig. 1B). To construct mimetic mutants, the *Eco*RI-*Eco*RI fragment amplified from pLF61-7D was cloned into the *Eco*RI site of vector Litmus39 (NEBiolabs) yielding L39-AS1-C, which served as quick change mutation template to engineer all four *P. anserina AS1* mutations using primers listed in Table S1. Mutated *Eco*RI-*Eco*RI fragments were then cloned into the unique *Eco*RI site of plasmid yEPU-*rps15*-ΔC. Resulting plasmids (*URA3, CEN*) were called yEPU-*rps15*-*AS1-1*, yEPU-*rps15-AS1-2*, yEPU-*rps15*-*AS1*-*4* and yEPU-*rps15*-*AS1*-*5*. A C-terminal truncated version of wild-type Sc*AS1* chimeric gene was constructed by cloning annealed oligonucleotides oS15-ΔC-F and oS15-ΔC-R into the *Eco*RI site of yEPU-*rps15*-ΔC. Oligonucleotides were designed to induce the replacement of Proline 138 (*P. anserina* numbering) by an ochre (TAA) stop codon so that the resulting plasmid yEPU-*rps15*-ΔCPro expresses a Sc*AS1ΔC* chimera truncated for the last 15 aas of *P. anserina* part of the WT version of chimeric uS19 r-protein.

A set of (*LEU2, CEN*) plasmids was then constructed by substituting in each yEPU plasmid the *Bgl*II-*Bgl*II *URA3* auxotrophic cassette by the *Bgl*II-*Bgl*II *LEU2* auxotrophic cassette PCR amplified from plasmid pFL36 using the primer pair FLK7w and FLK7c. Resulting plasmids were called yEPL-*RPS15*-*AS1*, yEPL-*rps15*-ΔCPro, yEPL-*rps15*-*AS1-1*, yEPL-*rps15*-*AS1-2*, yEPL-*rps15*-*AS1-4* and yEPL-*rps15*-*AS1-5*. They were used to transform yeast strain ME14-a9. All but yEPL-*rps15*-ΔCPro plasmid sustained growth on glucose, media in which the endogenous plasmid pFL38-*GAL::RPS15* will not express the essential *RPS15* copy. Strains were finally cured from this pFL38-*GAL::RPS15* (*URA3, CEN*) plasmid using 5-FOA and identity of the remaining plasmid (one of the yEPL set) was checked by PCR on colony and restriction enzyme diagnosis. Resulting strains were named according to the *S. cerevisiae*/*P. anserina* chimera they expressed: Sc*AS1* (wild-type chimera), Sc*AS1-1* (chimera mimicking the *AS1-1* change), Sc*AS1-2* (chimera mimicking *AS1-2* change), Sc*AS1-4* (chimera mimicking *AS1-4* change) and Sc*AS1-5* (chimera mimicking *AS1-5* change). All along DNA constructions, sequencing was performed on both strands to check cloned fragments and surrounding regions of cloning.

### Ribosomes and Polysomes purification

For *S. cerevisiae* strains, polysomes extractions were carried out starting from exponentially growing cultures in glucose rich medium at 30°C. Cells were treated with cycloheximide at a final concentration of 100 μg ml^-1^ immediately before harvesting to trap elongating ribosomes. Polysomes extractions were performed according to (Gregory *et al*., 2007).

For *P. anserina* strains, ribosomes and polysomes were isolated from mycelium grown on cellophane disks placed on M2 medium (27°C, two days). Mycelium was dried, frozen and placed into a nitrogen cooled Teflon shaking flask with four grinding stainless steel beads. Mycelium was crushed into fine powder using a Mikro-Dismembrator machine (Sartorius) and shaking at a frequency of 2600 per min for 90 seconds. For polysome extraction, collected mycelium was mixed with cycloheximide (100 μg ml^-1^) for ten minutes at 4°C prior drying, freezing and crushing. Ribosomes or polysomes were then cold-extracted in a volume of 1 ml for 150 mg of dried mycelium. Mycelium powder was re-suspended in extraction buffer [20 mM Tris-HCl (pH 8.0), 140 mM KCl, 1.5 mM MgCl_2_, 1 % Triton X-100, 0.5 mM DTT, 3 μl of RNasin ribonuclease inhibitor (Promega) *per* ml of extraction buffer]. For polysome extractions extra cycloheximide was added at a final concentration of 500 μg ml^-1^. Suspensions were first clarified by centrifugation at 4 000g for 5 min and the supernatant was further cleared at 12 000 g for 10 min. Extracts of equivalent of 10 OD_260_ units (approximately 400 μg of nucleic acids) were loaded on a 11 ml 7–50% sucrose gradient prepared in buffer [50 mM Tris HCl (pH 7.4), 50 mM NaCl, 1 mM DTT] for ribosome extracts and in buffer [50mM Tris Acetate (pH 7.5), 50 mM NH_4_Cl, 12 mM MgCl_2_, 1 mM DTT] for polysome extracts. Gradients were centrifuged at 38 000 rpm for 2 h 30 (polysomes) to 3 h 30 (ribosomes) at 4°C in a SW41 rotor.

### Nonsense codon readthrough assays

To quantify readthrough of nonsense codons, yeast strains Sc*AS1*, Sc*AS1-1*, Sc*AS1-2*, Sc*AS1-4* and Sc*AS1-5* were transformed with the control (pACU-TQ), UAG (pACU-UAG), UAA (pACU-UAA) or UGA (pACU-UGA) *LacZ-luc* reporter plasmid. Plasmid pACU-TQ carries the in-frame control, which allows production of 100% of β-galactosidase– luciferase fusion protein. All reporter plasmids are pAC derivative vectors (Stahl *et al*., 1995; Bidou *et al*., 2000) in which the *Bgl*II-*Bgl*II *LEU2* marker was replaced by the *Bgl*II-*Bgl*II *URA3* cassette by amplification from plasmid pFL38 using the primer pair FLK7w and FLK7c or by cloning of the *Bgl*II-*Bgl*II *URA3* cassette from plasmid pFL38. The 51 bp nonsense-containing sequence (GGG GAT CCC GCT AGC TGG CCA GCA GGA ACA CAA **STOP** CAA TTA CAG TGG CCA) is bordered by the natural 1024^th^ codon of the *lacZ* CDS and the natural 6^th^ codon of *luc* CDS (Fig. 5A). For each strain, the β-galactosidase and firefly luciferase activities were quantified in the same crude extracts as described (Stahl *et al*., 1995). The ratio of luciferase activity to β-galactosidase activity from the in-frame control construct (pACU-TQ) was taken as reference and termination codon readthrough (expressed in percent) was then calculated by dividing the luciferase/β-galactosidase ratio obtained from each nonsense construct by the same ratio obtained with the in-frame control construct.

## Acknowledgements

Thanks are due to Guillaume Canal, Raynad Cossard, Emmanuel Dassa, Agnés Delahodde, Michela Esposito, Laras Pitayu, Celine Roudier, Carole Sellem and Christelle Vasnier for their constructive review of the work and support. Tan-Trung Nguyen has been funded by a 3-year PhD Studentship from Paris-Sud University (Orsay, France). We also gratefully acknowledge additional support from the Franco-Vietnamese association from Paris-Sud University. This work was supported from the French Centre National de la Recherche Scientifique (CNRS) and the French Foundation FRM (INE20071110914 to Sylvie Hermann-Le Denmat).

## Author contributions

GS, VC and SHLD conceived and designed the study. TTN, VC, and MDC performed the *P. anserina* experiments. GS and SHLD performed the *S. cerevisiae* experiments. TTN, GS, MDC, VC and SHLD analyzed the data. VC and SHLD supervised the work. SHLD wrote the original draft. GS and SHLD made major contribution to the writing. All authors reviewed, edited and approved the final manuscript.

---

**Table S1.**
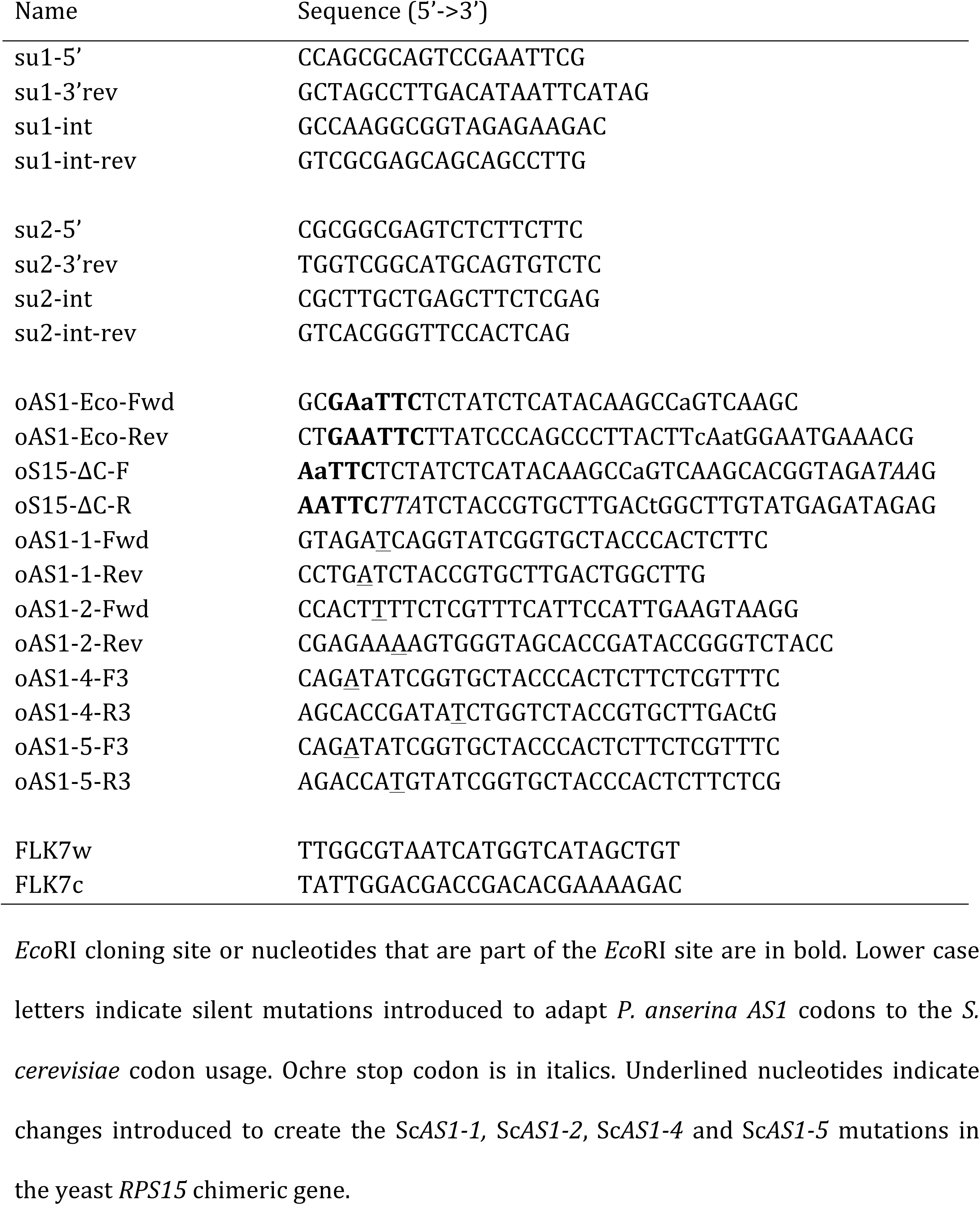
Oligonucleotides used in this study.

**Fig. S1.**
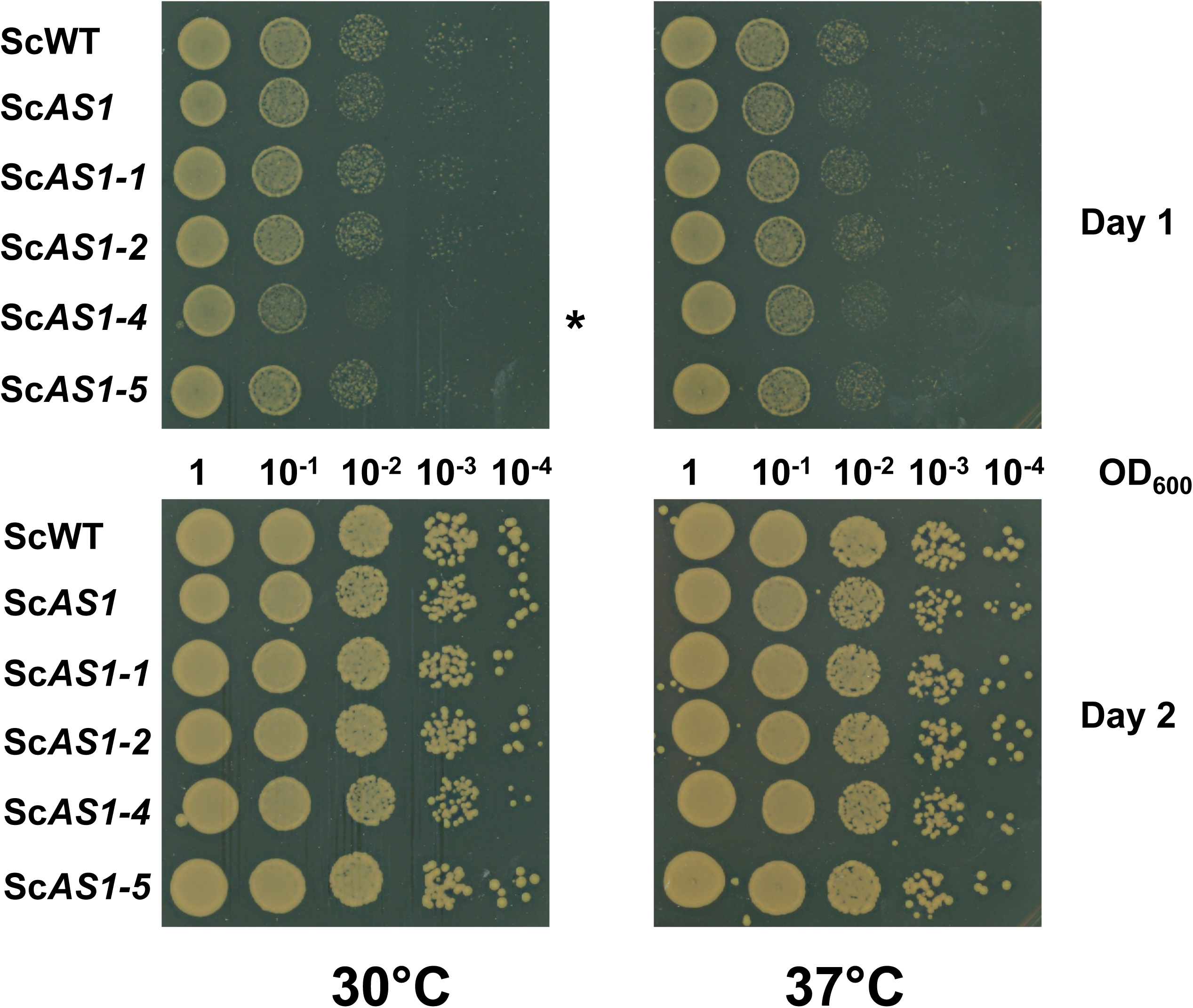
Growth delay of the *ScAS1-4* strain. Growth of equivalent 10-fold dilutions (indicating OD_600_) of the indicated yeast strains on glucose rich medium. Growth of the same plate was observed after one and two days of growth at the indicated temperature. The growth delay observed for the *ScAS1-4* strain at 30°C after one day of growth (*) fades at day 2 albeit individual colonies appear smaller than those of the reference *ScAS1* strain.

